# *Staphylococcus aureus* skin colonization is mediated by SasG lectin variation

**DOI:** 10.1101/2023.11.20.567970

**Authors:** Krista B. Mills, Joseph J. Maciag, Can Wang, John A. Crawford, Timothy J. Enroth, Klara C. Keim, Yves F. Dufrêne, D. Ashley Robinson, Paul D. Fey, Andrew B. Herr, Alexander R. Horswill

## Abstract

*Staphylococcus aureus* causes the majority of skin and soft tissue infections, but this pathogen only transiently colonizes healthy skin. However, this transient skin exposure enables *S. aureus* to transition to infection. Initial adhesion of *S. aureus* to skin corneocytes is mediated by surface protein G (SasG). Here, phylogenetic analyses reveal the presence of two major divergent SasG alleles in *S. aureus*, SasG-I and SasG-II. Structural analyses of SasG-II identified a unique non-aromatic arginine in the binding pocket of the lectin subdomain that mediates adhesion to corneocytes. Atomic force microscopy and corneocyte adhesion assays indicated SasG-II can bind to a broader variety of ligands than SasG-I. Glycosidase treatment resulted in different binding profiles between SasG-I and SasG-II on skin cells. Additionally, SasG-mediated adhesion was recapitulated using differentiated N/TERT keratinocytes. Our findings indicate that SasG-II has evolved to adhere to multiple ligands, conferring a distinct advantage to *S. aureus* during skin colonization.

## Introduction

The opportunistic pathogen *Staphylococcus aureus* is a common asymptomatic colonizer in humans but is also the predominant cause of skin and soft tissue infections ^1^. Approximately 20% of the human population are persistently colonized by *S. aureus,* most commonly in the anterior nares, while 30% are transiently colonized ^2,3^. *S. aureus* infections usually occur in individuals already colonized ^3,4^. Despite causing approximately 76% of skin and soft tissue infections ^5^, *S. aureus* only colonizes 5% or less of the skin of otherwise healthy adults ^6^. A trifecta of intact barrier, immune system, and commensal microbiota maintain skin homeostasis and keep *S. aureus* colonization rates low. However, dysbiosis or breaks in the skin barrier can lead to increased colonization levels and subsequent infection ^7–9^. When skin homeostasis is compromised, *S. aureus* can deploy virulence factors that enable immune evasion and tissue invasion, further exacerbating inflammation and disease ^10–12^. Identifying the mechanisms by which *S. aureus* colonizes healthy skin could open avenues for therapeutic development to prevent and treat infection in colonized individuals.

*S. aureus* skin colonization begins with initial adhesion, which is mediated by various cell wall-anchored (CWA) proteins ^11,13–15^. In particular, surface protein G (SasG) is known to be important in *S. aureus* adhesion to healthy human corneocytes on the skin ^15–17^, as well as adhesion to nasal epithelial cells and in biofilm formation ^18,19^. SasG is a large sortase-anchored protein that is part of the G5-E repeat family of adhesins ^11^. The structure of this protein consists of a N-terminal A domain encompassing an intrinsically disordered region and L-type lectin, a B domain with highly-conserved, serial B-repeats containing G5-repeat and E-spacer sub-domains ^11,16,18^, and a proline/glycine-rich stalk region extending to the C-terminus ^20^. SasG is orthologous to accumulation-associated protein (Aap) in *S. epidermidis*, and SasG homologs are expressed in other species of staphylococci as well ^15^. Similar to SasG, Aap has been found to be important in *S. epidermidis* adhesion to healthy human corneocytes ^15,16^.

Previous studies have elucidated the role of the A domain, specifically the L-type lectin subdomain, in Aap and SasG-mediated adhesion to corneocytes on healthy human skin ^15–17^. L-type lectins exhibit glycan-binding specificity that can significantly vary depending on the lectin^21^. While the specific ligand for Aap and SasG is still unknown, recent data has indicated the ligand is most likely a glycoprotein. Roy et al. ^15^ demonstrated that healthy human skin corneocytes treated with the glycosidases PNGase and O-Glycosidase significantly reduced *S. epidermidis* adhesion, suggesting these glycan linkages are important for adhesion to the host ligand. A glycan array performed by Maciag et al. ^16^ supported these data and found that the highest-affinity hits for the Aap lectin included N-linked glycans containing Gal-GlcNAc alternating repeats (poly-N-acetyllactosamine). Furthermore, Maciag et al. identified key aromatic residues within the glycan-binding pockets of the Aap and SasG lectins, which likely bind glycans through stacking interactions. These key aromatic residues are approximately in the same structural position, Y580 for Aap and W392 for SasG, and mutation of these residues to alanine abrogated binding of these lectins to N-acetyl-D-lactosamine and 3’-sialyl-N-acetyllactosamine as determined via isothermal titration calorimetry (ITC) ^16^. Similarly, pre-incubating corneocytes from healthy human skin with purified lectins from Aap and SasG significantly reduced adhesion of both *S. epidermidis* and *S. aureus*, while the mutated lectins did not affect adhesion ^16^, demonstrating that Aap and SasG bind to the ligand via these key residues in the lectin subdomain.

The ability of purified lectins from both Aap and SasG to bind lactosamine derivatives and reduce adhesion of both *S. epidermidis* and *S. aureus* to corneocytes suggests these species may compete for adhesion to the same ligand on healthy human skin. However, in a *S. epidermidis*-dominated skin environment ^22,23^ that is hostile to *S. aureus*, much has yet to be elucidated as to what makes SasG unique and able to establish a colonization niche. Here, we use phylogenetic analyses to demonstrate the presence of two divergent allelic types of SasG, with each type represented by full-length and truncated forms, within *S. aureus*. We demonstrate that SasG-II is unique and may bind to multiple ligands, based on evidence that includes the SasG-II lectin structure and a lectin alignment comparison to SasG-I, adhesion studies on corneocytes from healthy human skin and immortalized N/TERT keratinocytes, and nanoimaging of corneocytes using SasG-I and SasG-II *S. aureus* single cell probes.

## Results

### The SasG A domain is variable in *S. aureus*

Variability in the repertoire of sortase-attached adhesins is common on the surface of *S. aureus* ^24^. SasG in particular has been noted as having significant strain-level diversity in gene (*sasG*) presence, expression level, and function ^18,25^. We realized that *S. aureus* strains expressing full-length SasG, such as well-known strains COL and 502a in clonal complex 8 (CC8) and CC5, and MW2 in CC1, have considerable divergence in the A domain sequence. Considering the SasG A domain has been recently linked to corneocyte adhesion ^15,17^, we reasoned that the sequence differences could impact *S. aureus* skin colonization.

Full-length SasG from MRSA strain COL has a lectin domain that is structurally similar to that of *S. epidermidis* Aap, as we recently reported ^16^. We named this form of SasG as Type I or “SasG-I” hereafter. In contrast, the A domain lectin in MRSA strain MW2 is only 67.2% identical at the protein level. We named this form of SasG as Type II or "SasG-II" hereafter. The rest of the protein content in both full-length SasG forms is fairly similar with the B-repeat region containing G5 and E spacers and a C-terminal wall/membrane spanning region with a LPKTG sortase motif (**Fig. S1A**). In this initial analysis, we also determined that some strains encoding SasG-I had an intact A domain and a frameshift mutation at the start of the B-domain. This includes strains such as N315 ^26^ and LAC ^19^ that belong to USA100-related and USA300 lineages, respectively. We named this mutated form of SasG-I as “truncated SasG-I”. Some strains with SasG-II also encode a truncated protein; however, this form occurs less frequently among *S. aureus* strains and was not investigated further.

Both the Aap and SasG-I lectins have a key active site aromatic residue that has been linked to glycan binding and corneocyte adhesion, which is Y580 in Aap and W392 in SasG-I ^16^. Based on an alignment of the SasG-I and SasG-II lectins, this aromatic residue is missing in SasG-II and is replaced with non-aromatics (**Fig. S1B**). Other secondary structure elements are similar, as expected given that both adopt β-sandwich L-type lectin folds. Based on our previous analysis of the SasG-I and Aap lectin structures ^16^, most of the key residues in the vicinity of the glycan binding site (shortly after β17) are conserved between SasG-I and SasG-II, with one important exception, as described below.

### Phylogenetic analyses reveal the depth of SasG variation

Building on our preliminary observations, we further investigated SasG variation using a phylogenetically diverse set of *S. aureus* isolates for which whole genome sequences were available ^27^. These 574 isolates belonged to 39 clonal complexes (**Fig. 1A**). Among these isolates, SasG was represented by three approximately evenly distributed groups: 191 full-length SasG, 194 truncated SasG, and 189 missing SasG. Phylogenetic analysis of the aligned full-length SasG sequences confirmed that the species has two divergent allelic types with 99 inferred amino acid substitutions between them (**Fig. 1C**, long branch separating the two types). Among the full-length SasG sequences, there were 91 unique SasG sequences with 11 distinct amino acid lengths of sequence that could be attributed to the B-repeat region. The individual B-repeats were exactly 128 amino acids in length, and 11 differently sized repeat arrays were detected among the full-length SasG sequences. The distribution of B-repeats among the two full-length SasG types was significantly different (χ^2^=57.3, df=10, *P*<0.0001), with a more even number of B-repeats among the Type II isolates compared to the Type I isolates (**Fig. 1B**).

**Figure 1.**
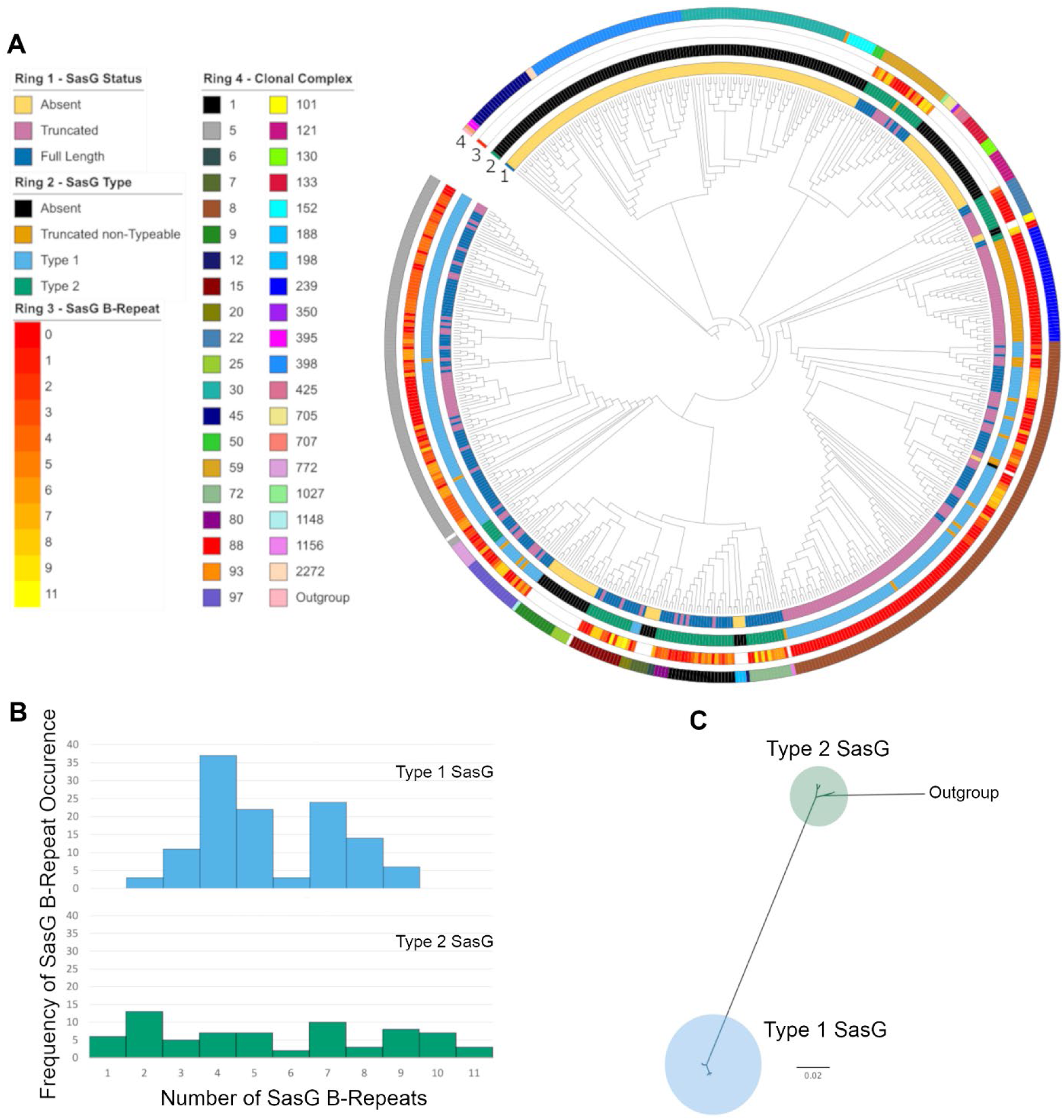
SasG Variation in the context of *S. aureus* phylogenetic diversity. (**A**) Maximum likelihood phylogenetic tree of 574 *S. aureus* isolates based on their proteome. For each isolate, SasG status is mapped to ring 1, SasG type (defined in Panel B) is mapped to ring 2, SasG B-repeat number is mapped to ring 3, and the clonal complex of the isolate is mapped to ring 4. (**B**) Histograms of SasG B-repeats from both SasG allelic types. (**C**) Maximum likelihood phylogenetic tree of SasG from 191 aligned full-length sequences identifies two SasG allelic types.

Different isolates of a given clonal complex always had the same SasG type. However, at least four changes of SasG types are inferred based on parsimony analysis of the type distribution on the *S. aureus* phylogeny. The two SasG types correlate perfectly with amino acid polymorphisms in the L-type lectin binding region: W392 for SasG-I, and R394 for SasG-II. Since polymorphisms in these alignment positions correlated perfectly with the full-length SasG types, these 2 amino acids were used to type the truncated SasG sequences. This typing allowed for an additional parsimony analysis that identified a minimum of 27 changes between a predicted functional SasG (full-length or typed-truncated sequence) and a predicted nonfunctional SasG (missing or untyped-truncated sequence). The clonal complexes CC5 and CC8, which are clinically important ^28–30^, both encode SasG-I but CC5 has mostly full-length forms and CC8 has mostly truncated forms. SasG-II is encoded by CC1 (such as MRSA strain MW2), as well as CC15, CC22, CC59, CC72, and others. SasG-II in these CCs are mostly full-length, although some strains do rarely harbor truncated SasG-II.

### The SasG-II lectin contains a unique non-aromatic residue in the glycan binding pocket

The crystal structure of SasG-II lectin was solved at 1.88 Å, revealing an overall architecture that is nearly identical to the L-type lectin folds of SasG-I, Aap, and Pls ^16,31^ (**Fig. 2A**). Similar to these related lectin domains, SasG-II has an atypical *trans* conformation of the central D241 residue, a structural Ca^2+^ ion, and three relatively long loops at the top of the domain. However, both Aap and SasG-I lectins feature a sharp bend in the main chain after strand β17 that positions an aromatic residue at the base of the glycan binding pocket (i.e., Aap Y580 and SasG-I W392) (**Fig. 2B**). In contrast, SasG-II fails to adopt the sharp bend at the end of β17, which therefore positions the sidechain of R394 in approximately the same position as the SasG-II main chain after the bend. The arginine at this position is actually conserved between SasG-I and SasG-II, but the sharp bend in the main chain of SasG-I causes the equivalent R391 to extend toward the top of the lectin domain, far from the binding pocket, and positions W392 within the binding pocket instead (**Fig. 2C**). Likewise, the Q395 residue in SasG-II (corresponding to W392 in SasG-I) points in the opposite direction to R394 in a region of the protein that is highly solvent-exposed (**Fig. 2C**). It seems likely that SasG-I and Aap adopt the sharp bend in the main chain after β17 to avoid exposure of the equivalent aromatic residues (W392 or Y580) to solvent, which would be entropically unfavorable. Instead, the main chain bend orients those aromatic residues toward the binding site pocket region, which is more protected from solvent. As a result of these structural differences, SasG-II forms a surface pocket similar to the other lectins’ glycan binding sites but it does not contain an aromatic residue that could form a stacking interaction with a glycan ligand (**Fig. 2D**).

**Figure 2.**
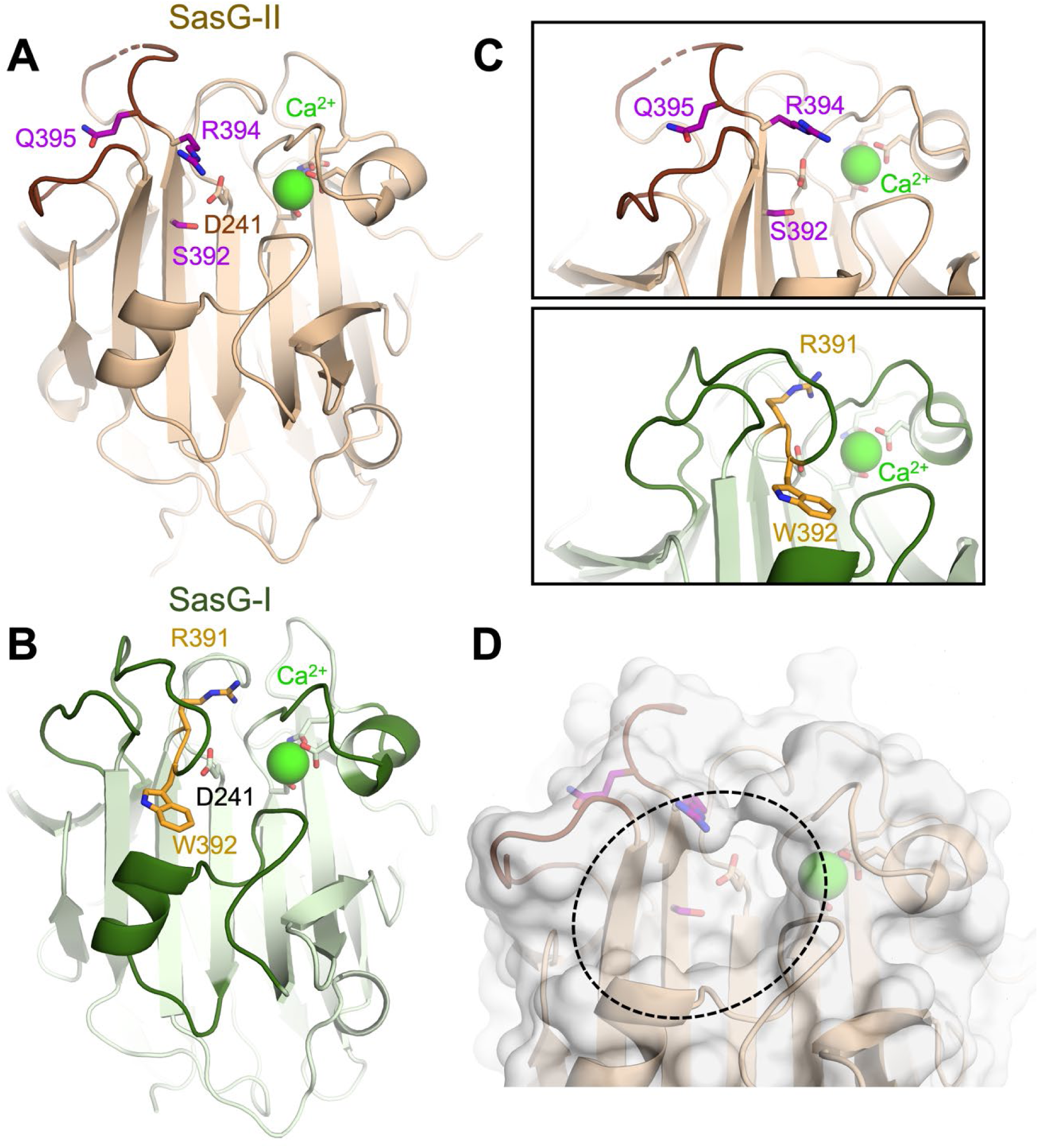
The SasG-II lectin contains a unique non-aromatic residue in the glycan binding pocket. (**A**) Crystal structure of the SasG-II lectin showing the structural Ca^2+^ ion, the conserved central D241 residue that adopts an atypical *trans* conformation, and the side chains of S392, R394, and Q395 near the end of β17. (**B**) Comparative view of SasG-II in the same orientation. Residues R391 and W392 are analogous to R394 and Q395 in SasG-II; note the distinct positioning of corresponding residues W392 (SasG-II) and Q395 (SasG-I). (**C**) Close-up view of the region near the end of β17, rotated by approximately 45° from panels A and B. Note the sharp bend of the main chain near R391 in SasG-I that is not observed in SasG-II. (**D**) Surface view of SasG-II showing the putative binding pocket lacking an aromatic residue at its base.

### Multiparametric nanoimaging using single bacterial probes indicates SasG-II binds a broader variety of ligands than SasG-I

To compare SasG-I and SasG-II-mediated adhesion at the single molecule level and further explore the possibility of SasG-II binding multiple ligands, we determined the strength of adhesion between a single living bacterial cell and human corneocytes using multiparametric atomic force microscopy (AFM) imaging ^32^. Bacterial cells not expressing SasG (*S. carnosus*-pALC2073 EV) did not show any adhesion (**Fig. 3**), similar to a colloidal probe (**Fig. 3 and Fig. S2**). Bacterial cells expressing SasG-II (*S. carnosus-sasG*_MW2_) showed strong adhesion forces ranging from 500 to 5,000 pN, with a most frequently observed force of around 1,000 pN, that were densely and widely distributed across the corneocyte surface (**Fig. 3 and Fig. S3**). It is likely that the 1,000 pN force corresponds to a single interaction, while larger forces represent multiple bonds, possibly of a different molecular nature. The detection frequency of 27% demonstrates that the SasG-II ligand(s) are present at high density. SasG-I-expressing cells also showed many adhesion forces around 1,000 pN, yet with a much lower detection frequency of approximately 7% (**Fig. 3 and Fig. S4**). This indicates that both adhesins strongly bind their skin ligands, yet SasG-II seems to bind a broader variety of ligands.

**Figure 3.**
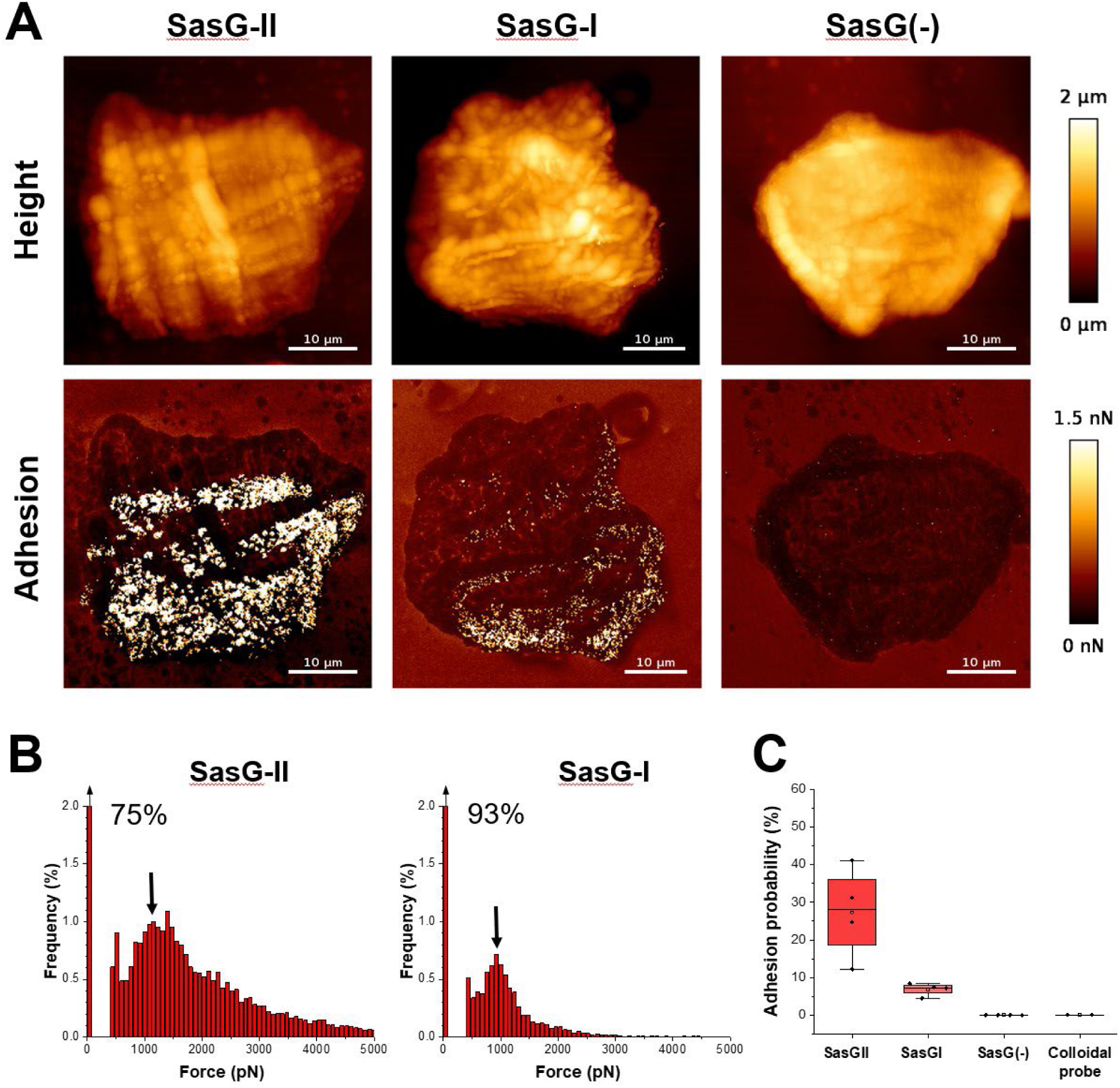
Multiparametric nanoimaging using single bacterial probes indicates SasG-II binds a broader variety of ligands than SasG-I. (**A**) Height images (top) and adhesion images (bottom) of corneocytes recorded in PBS using a SasG-II, SasG-I, or EV (SasG[-]) cell probe. See also Figure S2-S4. (**B**) Histograms of adhesion forces registered on whole corneocytes (total of n = 9,590 curves for one representative SasG-II probe; n = 2,532 curves for one representative SasGI probe). The arrow at the top left of the histograms stands for the non-adhesive events. (**C**) Box plot comparing adhesion probabilities for SasG-II (n = 4 from 3 independent bacterial cultures), SasG-I (n = 4 from 2 independent bacterial cultures), SasG(-) (n = 4 from 2 independent bacterial cultures) or colloidal (n = 2) probes. For more data, see Figures S1, S2, and S3.

### Purified glycans do not affect SasG-II-mediated adhesion to corneocytes

As we have shown previously, the Aap and SasG-I lectins were both able to bind purified N-acetyl-D-lactosamine via ITC ^16^. Using ITC, we investigated whether SasG-II would also bind N-acetyl-D-lactosamine, and found that the SasG-II lectin did not bind (**Fig. S5A**). This was functionally validated in a corneocyte adhesion assay, where pre-incubation with a serial 2-fold dilution of N-acetyl-D-lactosamine ranging from 1000 µM to 62.5 µM did not affect SasG-II-expressing *S. carnosus-sasG*_MW2_ adhesion to corneocytes (**Fig. S5B and S5C**). Similarly, pre-incubation with a serial 2-fold dilution of 3’-sialyl-N-acetyllactosamine ranging from 100 µM to 6.25 µM (10-fold lower dilutions than was used with N-acetyl-D-lactosamine) did not affect SasG-II-expressing *S. carnosus-sasG*_MW2_ adhesion to corneocytes (**Fig. S5D and S5E**). In contrast, adhesion of SasG-I-expressing *S. carnosus*/pALC2073-*sasG*_COL_ was strongly inhibited by 125 µM N-acetyl-D-lactosamine or by 12.5 µM 3’-sialyl-N-acetyllactosamine ^16^. These data collectively suggest that SasG-II is unique from Aap and SasG-I, in that it may bind distinct glycan ligands on the corneocyte receptor, or perhaps that it binds the protein portion of the receptor without engaging the glycan moieties.

### SasG-I and -II mediated adhesion to corneocytes shows differential responses upon treatment with glycosidases

To narrow down possible ligand configurations for SasG-I and SasG-II and further investigate binding differences between the two types, corneocytes were pre-incubated with glycosidases. Protein glycosylation has been postulated to be important for epidermal differentiation, as well as desquamation, hydration, and adhesion/cohesion of the stratum corneum ^33,34^. The most common types of glycans found on healthy human skin are variations of complex N-linked glycans and mucin-like core 1 and core 2 O-linked glycans ^34–37^. Seven different glycosidases were tested, including endoglycosidases and exoglycosidases. PNGase F (cleaves between the innermost GlcNAc and asparagine residues of high-mannose, hybrid, and complex oligosaccharides), O-Glycosidase (catalyzes removal of Core 1 and 3 O-linked disaccharides from glycoproteins), α1-2,3,6 Mannosidase (catalyzes the hydrolysis of terminal, non-reducing α1-2, α1-3 and α1-6 linked mannose residues from oligosaccharides), α1-3,4 Fucosidase (catalyzes the hydrolysis of terminal, non-reducing α1-3 and α1-4 linked fucose residues from oligosaccharides and glycoproteins), β-*N*-Acetylglucosaminidase S (catalyzes the hydrolysis of terminal, non-reducing β-N-Acetylglucosamine residues from oligosaccharides), β1-4 Galactosidase S (catalyzes the hydrolysis of terminal, non-reducing β1- 4 linked galactose residues from oligosaccharides), and α2-3,6,8 Neuraminidase (catalyzes the hydrolysis of α2-3, α2-6, and α2-8 linked sialic acid residues from glycoproteins and oligosaccharides).

We predicted that SasG-I-mediated adhesion would be similar to our previous results with Aap ^15^, and would be reduced upon glycosidase treatment with the endoglycosidases PNGase F and O-Glycosidase, as well as α2-3,6,8 Neuraminidase, and the exoglycosidases α1-3,4 Fucosidase and β1-4 Galactosidase S. Indeed, pre-incubation with all five of these glycosidases resulted in significantly reduced adhesion of *S. carnosus-sasG*_COL_ compared to buffer controls, while the exoglycosidases α1-2,3,6 Mannosidase and β-*N*- Acetylglucosaminidase S did not affect adhesion (**Fig. 4A-G**). This suggests that linkages between the innermost GlcNAc and asparagine residues of oligosaccharides, as well as Core 1 and 3 *O*-linkages are important in SasG-I mediated adhesion, and that 5-*N*-acetylneuraminic acid, fucose, and galactose may be important in the SasG-I ligand configuration.

**Figure 4.**
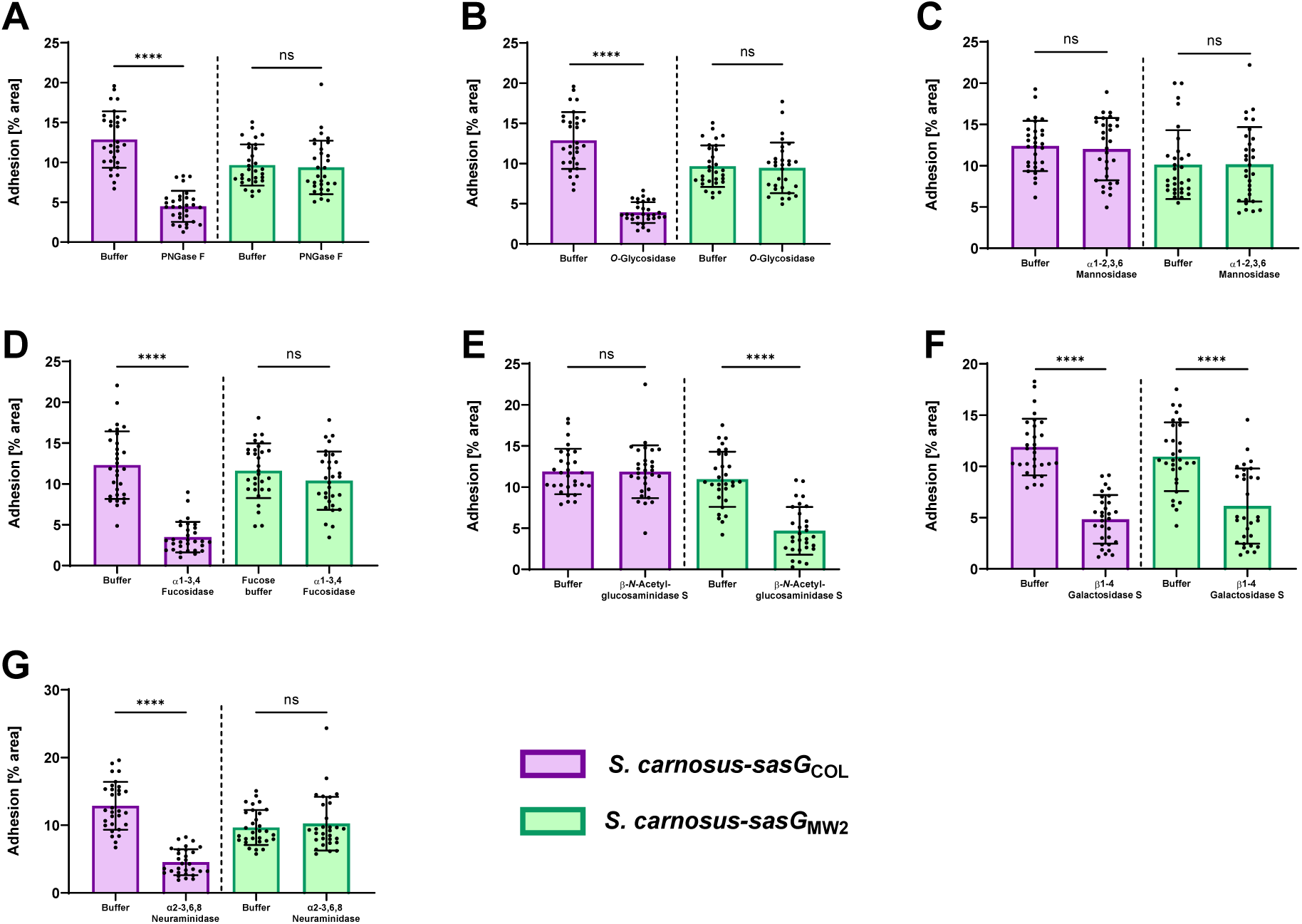
SasG-I and -II mediated adhesion to corneocytes shows differential responses upon treatment with glycosidases. *S. carnosus-sasG*_COL_ (SasG-I) and *S. carnosus-sasG*_MW2_ (SasG-II) were tested for adhesion to corneocytes following pre-incubation with (**A**) PNGase F, (**B**) O-Glycosidase, (**C**) α1-2,3,6 Mannosidase, (**D**) α1-3,4 Fucosidase, (**E**) β-*N*- Acetylglucosaminidase S, (**F**) β1-4 Galactosidase S, and (**G**) α2-3,6,8 Neuraminidase. The percent area of adhesion in 10 images from three independent experiments (n = 30) was measured with Fiji ImageJ and analyzed in GraphPad Prism. Statistical significance was analyzed using the unpaired t-test or non-parametric Mann-Whitney test for data with non-normal distribution (\*\*\*\**P*<0.0001).

Considering the inability of the SasG-II lectin to bind lactosamine (**Fig. S5A**), we predicted that SasG-II may bind different terminal sugar residues on corneocytes than SasG-I. Using the same corneocyte adhesion assay, only pre-incubation with β-*N*- Acetylglucosaminidase S and β1-4 Galactosidase S resulted in a reduction of *S. carnosus-sasG*_MW2_ adhesion (**Fig. 4A-G**). This indicates that terminal β-*N*-acetylglucosamine and galactose may be important in the SasG-II ligand configuration. The only glycosidase that resulted in both reduced SasG-I and SasG-II adhesion was β1-4 Galactosidase S, suggesting that galactose may be important in the configuration of a shared ligand between SasG-I and SasG-II. These results further suggest that SasG-II binds a different type of glycan ligand than SasG-I on the corneocyte receptor.

### SasG-II-mediated adhesion is mediated by the lectin subdomain and may bind the same ligand as Aap and SasG-I

To investigate the possibility of a shared ligand between SasG-I and SasG-II, we sought to determine if purified lectins from Aap, SasG-I, and SasG-II could cross-inhibit SasG-II-mediated adhesion to corneocytes. Pre-incubating/blocking corneocytes from healthy human skin with 5 µM purified lectin domains from Aap, SasG-I, and SasG-II significantly reduced SasG-II-expressing MRSA MW2 adhesion, while purified lectins with the key residues mutated–Aap ΔY580A and SasG-I ΔW392A–did not affect adhesion (**Fig. 5A and 5B**). We then investigated whether this would hold true in the model surrogate organism *S. carnosus*, which does not natively adhere to corneocytes ^17^. Expressing SasG in *S. carnosus* acts as an ideal model for testing protein-specific adhesion since there are no other adhesins that could affect corneocyte binding. Similar to *S. aureus*, adhesion of *S. carnosus* expressing SasG-II on plasmid pALC2073 was significantly reduced when pre-incubated/blocked with purified lectins from Aap and SasG-I, but was not affected by pre-incubation with Aap ΔY580A or SasG-I ΔW392A, validating the findings in *S. aureus* (**Fig. 5C and 5D**). These data suggest that SasG-II may bind to the same host ligand as SasG-I and Aap. Alternatively, SasG-II could be binding elsewhere on the corneocyte receptor, sterically blocking SasG-I and Aap from binding.

**Figure 5.**
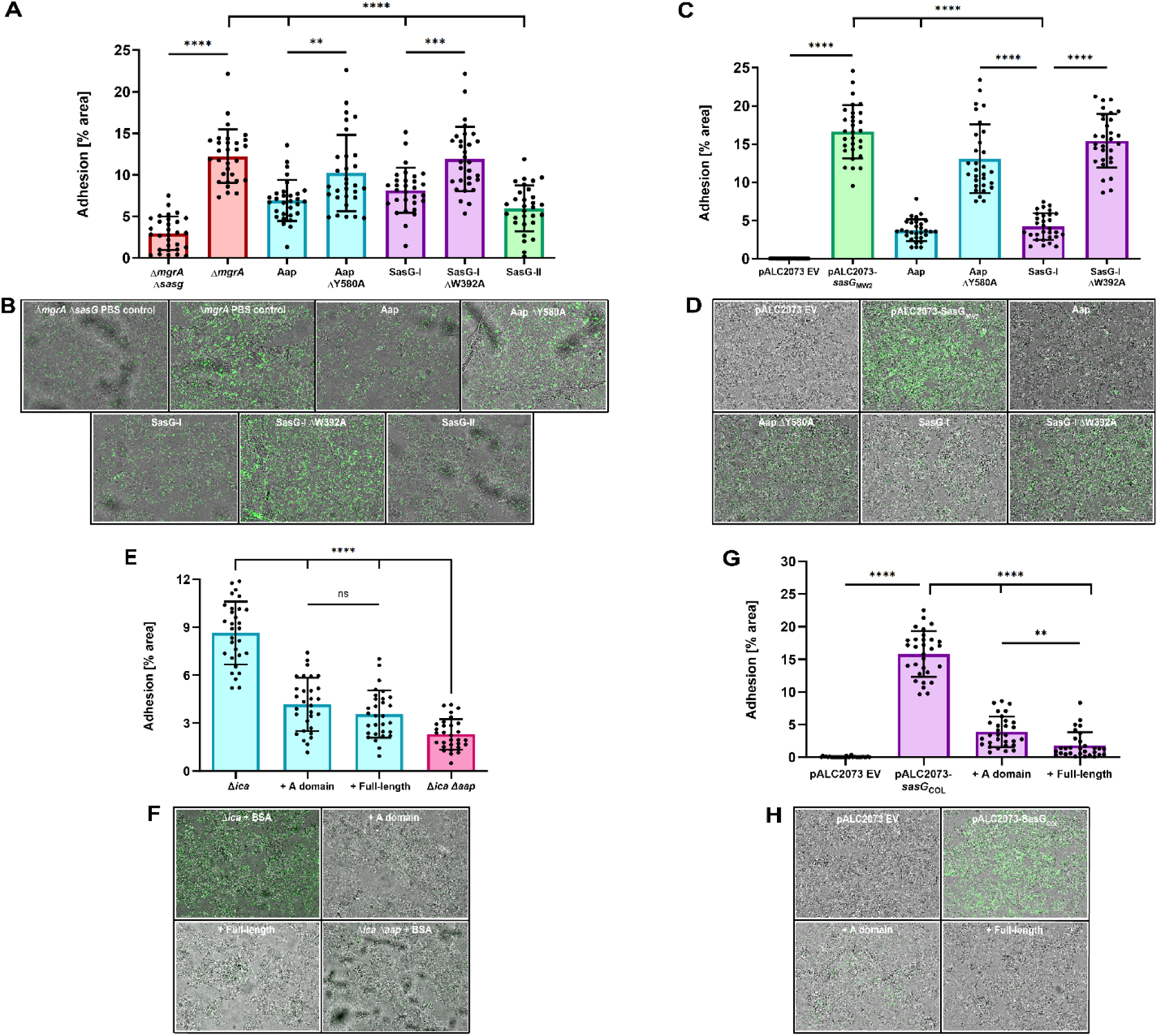
SasG-II-mediated adhesion is mediated by the lectin subdomain and may bind the same ligand as Aap and SasG-I. (**A**) MRSA MW2 (SasG-II) was tested for adhesion to healthy human corneocytes after pre-incubation/blocking with 5 µM of purified lectins. MRSA MW2 Δ*sasG* was used as a negative control strain. \*\**P*=0.0019. (**C**) SasG-II-expressing *S. carnosus-sasG*_MW2_ was tested for adhesion to healthy human corneocytes after pre-incubation/blocking with 5 µM of purified lectins. *S. carnosus*-pALC2073 EV was used as a negative control strain. *S. epidermidis* Δ*ica* was tested for adhesion to corneocytes following pre-incubation/blocking with 100 µg/mL of purified full-length or A-domain SasG-II. *S. epidermidis* Δ*ica* Δ*aap* was used as a negative control strain. (**G**) SasG-I-expressing *S. carnosus-sasG*_COL_ was tested for adhesion to healthy human corneocytes after pre-incubation/blocking with 100 µg/mL of purified full-length or A-domain SasG-II. *S. carnosus*- pALC2073 EV was used as a negative control strain. \*\**P*=0.0091. (**B, D, F, H**) Representative bright-field (representing corneocytes) and green-channel (representing GFP-expressing bacteria) overlay microscopy images of experimental groups tested in Panels A, C, E, and G, respectively. (**All panels**) The percent area of adhesion in 10 images from three independent experiments (n = 30) was measured with Fiji ImageJ and analyzed in GraphPad Prism. Statistical significance was analyzed using ordinary one-way ANOVA (\*\*\**P*=0.0002; \*\*\*\**P*<0.0001).

We then investigated if purified SasG-II could cross-inhibit SasG-I and Aap-mediated adhesion to corneocytes. Purification of Type II full-length and A-domain SasG from MRSA MW2 was characterized previously ^19^. Pre-incubation/blocking with 100 µg/mL purified A domain and full-length SasG-II significantly reduced *S. epidermidis* adhesion to corneocytes (**Fig. 5E and 5F**). Likewise, pre-incubation/blocking with 100 µg/mL purified A domain and full-length SasG-II significantly reduced SasG-I-expressing *S. carnosus-sasG*_COL_ adhesion to corneocytes (**Fig. 5G and 5H**). These findings suggest that the SasG-II mode of binding to the corneocyte receptor enables competition with staphylococci expressing Aap or SasG-I.

### SasG-I and SasG-II-mediated adhesion to differentiated N/TERT keratinocytes following treatment with glycosidases suggests complex N-linked glycans and core 2 O-glycans may be important for SasG-I and SasG-II binding

Firstly, to validate the adhesion data seen on corneocytes and explore whether SasG could be utilized for invasion, we utilized a live, immortalized, N/TERT-2G keratinocyte cell culture model. N/TERT keratinocytes can undergo terminal differentiation, allowing for the presence of multiple keratinocyte layers with a desquamated stratum corneum. Terminally differentiated N/TERT keratinocytes with a stratum corneum were incubated with *S. carnosus-*pALC2073 EV (negative control), SasG-I-expressing *S. carnosus*-*sasG*_COL_, SasG-II-expressing *S. carnosus*-*sasG*_MW2_, and SasG-II-expressing *S. carnosus* with the A domain deleted (*S. carnosus*-*sasG*_MW2ΔA_) ^17^. Similar to what was seen in the corneocyte adhesion assays, both SasG-I and SasG-II-expressing *S. carnosus* strains adhered significantly more to differentiated N/TERTs than the EV and SasG-II A domain mutant controls (**Fig. 6A and 6B**). To determine if SasG could be used not only in initial adhesion to the stratum corneum but also for adherence to the more basal, actively dividing keratinocyte layers, these same strains were then tested for adhesion to an undifferentiated monolayer of N/TERT keratinocytes. Conversely, none of the strains adhered differently in a statistically significant manner (**Fig. 6C and 6D**). The difference in adhesion for SasG-I-expressing *S. carnosus*- *sasG*_COL_ and SasG-II-expressing *S. carnosus*-*sasG*_MW2_ on differentiated vs undifferentiated N/TERT keratinocytes is illustrated in **Fig. 6E**, where much lower adhesion is seen for both strains in undifferentiated N/TERT keratinocytes than differentiated N/TERT keratinocytes containing a stratum corneum. These data indicate SasG is utilized by *S. aureus* for initial adhesion to the outermost layer of the skin to establish a colonization niche, but not to infiltrate deeper layers of the epidermis.

**Figure 6.**
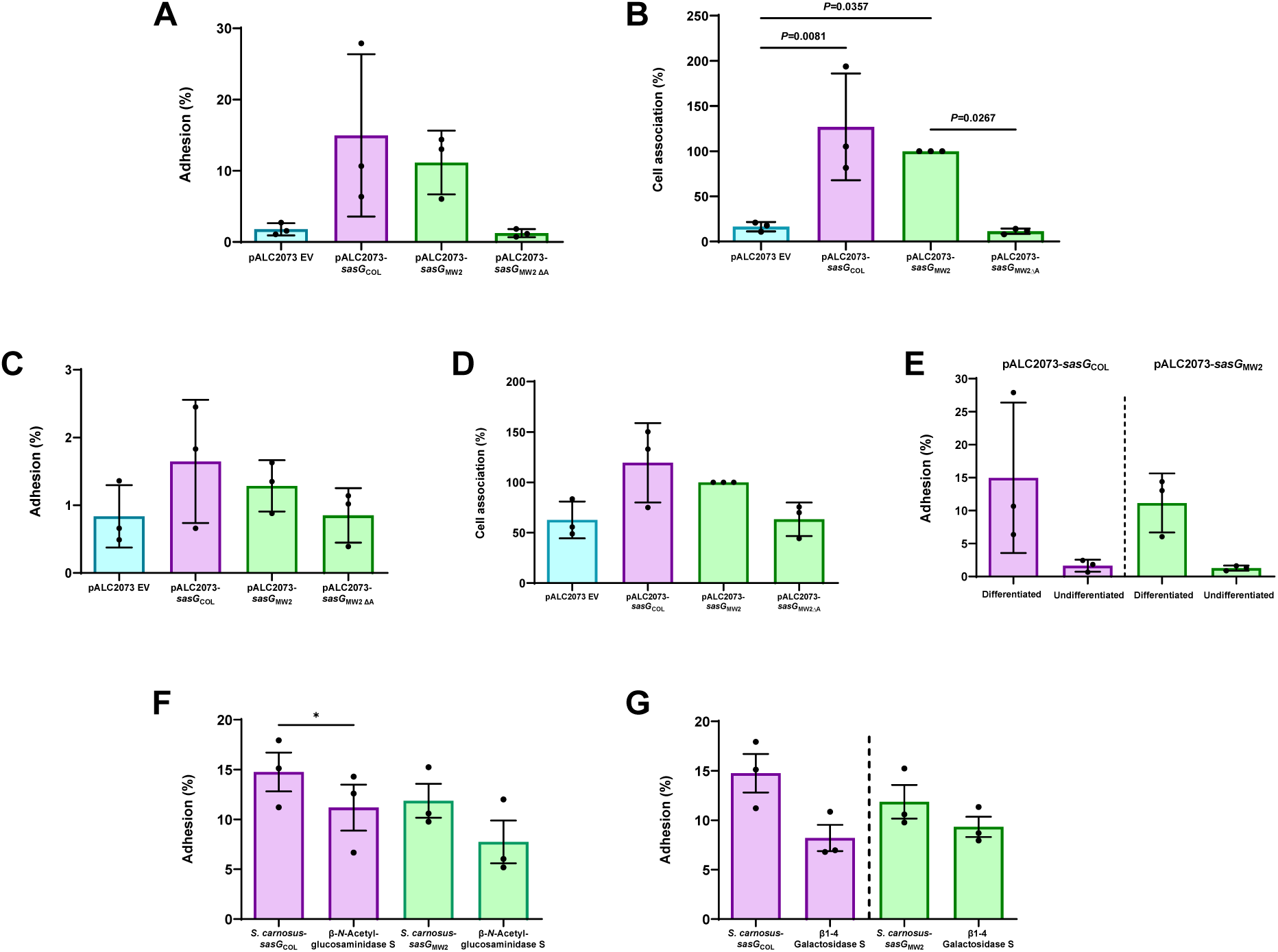
SasG-I and SasG-II-mediated adhesion to differentiated N/TERT keratinocytes following treatment with glycosidases suggests complex N-linked glycans and core 2 O-glycans may be important for SasG-I and SasG-II binding. *S. carnosus*-pALC2073 EV, SasG-I-expressing *S. carnosus-sasG*_COL_ and SasG-II-expressing *S. carnosus-sasG*_MW2_, and SasG-II-expressing *S. carnosus* with the A-domain deleted (*S. carnosus-sasG*_MW2ΔA_) at an MOI of 5 were tested for adhesion to either differentiated (**A and B**) or undifferentiated (**C and D**) N/TERT keratinocytes. (**A**) Adhesion to terminally differentiated cells as shown by overall percent adhesion. (**B**) Adhesion to terminally differentiated cells as shown by percent cell association to pALC2073-*sasG*_MW2_ input inoculum. Both SasG-expressing strains adhered more to differentiated cells than the EV and A domain mutant controls. (**C**) Adhesion to a monolayer of undifferentiated cells as shown by overall percent adhesion. (**D**) Adhesion to a monolayer of undifferentiated cells as shown by percent cell association to pALC2073-*sasG*_MW2_ input inoculum. There were no significant differences in adhesion between the EV and A domain mutant controls and the SasG-expressing strains. (**A-D**) The CFU/mL of three independent experiments (n = 3) were calculated and analyzed for statistical significance in GraphPad Prism using ordinary one-way ANOVA. (**E**) Data from panels A and C displaying differences in adhesion between differentiated and undifferentiated N/TERT keratinocytes for SasG-I-expressing *S. carnosus-sasG*_COL_ and SasG-II-expressing *S. carnosus-sasG*_MW2_. Both strains adhered well to differentiated N/TERT keratinocytes, and did not adhere well to undifferentiated N/TERT keratinocytes. No statistical analyses were performed for Panel E. SasG-I-expressing *S. carnosus-sasG*_COL_ and SasG-II-expressing *S. carnosus-sasG*_MW2_ were tested for overall percent adhesion to differentiated N/TERT keratinocytes following treatment with (**F**) β-*N*- Acetylglucosaminidase S and (**G**) β1-4 Galactosidase S. β1-4 Galactosidase S reduced adhesion of both strains, while β-*N*-Acetylglucosaminidase S resulted in a greater reduction in adhesion of *S. carnosus-sasG*_MW2_. The CFU/mL of three independent experiments (n = 3) were calculated and analyzed for statistical significance in GraphPad Prism using an unpaired t-test. \**P*=0.0500.

With the understanding that the ligand(s) to SasG exist on the stratum corneum, we then investigated whether the glycosidase treatment on corneocytes would reveal similar glycan ligand targets on live cells. *S. carnosus*-*sasG*_MW2_ and *S. carnosus*-*sasG*_COL_ were tested for adhesion to differentiated N/TERT keratinocytes following pre-incubation with glycosidases β-*N*- Acetylglucosaminidase S and β1-4 Galactosidase S. Similar to the corneocyte data, treatment with these two glycosidases reduced *S. carnosus*-*sasG*_MW2_ adhesion to differentiated N/TERT keratinocytes. Though not statistically significant, β-*N*-Acetylglucosaminidase S exhibited a reduced adhesion trend for *S. carnosus*-*sasG*_MW2_, and this reduction was greater than what was seen for *S. carnosus*-*sasG*_COL_ (**Fig. 6F**). β1-4 Galactosidase S reduced adhesion of both *S. carnosus*-*sasG*_COL_ and *S. carnosus*-*sasG*_MW2_, and resulted in a statistically significant reduction for *S. carnosus*-*sasG*_COL_ (**Fig. 6G**). These data confirm what was observed on corneocytes that galactose may be important in the configuration of a shared ligand between SasG-I and SasG-II, and that β-*N*-Acetylglucosamine may be uniquely important to a separate SasG-II ligand configuration. Terminal β1-4-linked galactose and β-*N*-acetylglucosamine are found within hybrid N-linked glycans ^34^, consistent with our previous observations that Aap binds to N-glycans from the glycan array and that PNGase F abrogates adhesion mediated via SasG-I or Aap ^16^. However, terminal β1-4-linked galactose and β-*N*-acetylglucosamine are also both found in core 2 mucin-type O-glycans ^38^. SasG-II may interact with this type of O-glycan structure, which would be consistent with the data that SasG-II-mediated adhesion is insensitive to treatment with PNGase F or O-glycosidase (O-glycosidase can cleave core 1 or core 3, but not core 2, O-glycans) (**Fig. 4**) ^39^. The glycome of healthy skin prominently features numerous N-glycoforms containing lactosamine or sialyllactosamine as well as core 2 O-glycans ^35,36^.

## Discussion

With over 50% of bacterial infections becoming resistant to treatment ^40^, identifying novel ways to prevent infection has never been more imperative. One of the human body’s first defense mechanisms against external hazards is the human skin ^41^. *S. aureus* plays an important role in the skin environment, causing the majority of skin and soft tissue infections ^5^ and contributing to morbidity in diseases such as atopic dermatitis ^42–44^. Recent studies have emphasized the importance of CWA adhesins in establishing colonization of human skin corneocytes, which comprise the outermost layer of the skin ^13–18^. One of these adhesins– SasG–is a multifactorial protein that shares structural and functional similarity to *S. epidermidis* Aap and has been characterized to be important in adhesion to corneocytes from healthy human skin ^15–17^. In this study, we identified variation in SasG via phylogenetic analyses, and investigated how this variation affects adhesion to healthy human skin using a multifaceted approach.

Phylogenetic analyses using a curated data set of the complete genomes from 574 *S. aureus* isolates revealed approximately a third of strains express a full-length form and another third express a truncated form of SasG. SasG-I is more than twice as common as SasG-II across the diverse clonal complexes of *S. aureus* examined here. Lastly, approximately a third of *S. aureus* strains do not encode SasG. Fibronectin binding protein B was recently found to important in healthy corneocyte interactions and might explain the ability of *S. aureus* to bind without SasG ^13^. SasG-II occurs in some of the earlier diverging clonal complexes, but also occurs in more derived clonal complexes likely due to a historical recombination event; it is not likely that the two allelic types could independently originate multiple times through mutation alone due to the large number of specific differences that define the two types. The observation that both allelic types of SasG exist in full-length and truncated forms in currently circulating strains, may suggest that retaining this molecular and functional variation is beneficial to *S. aureus* as a species. Thus, SasG may be a new candidate of balancing selection in *S. aureus*and needs to be further investigated ^45^.

The SasG-II lectin shares 67.2% sequence identity with the SasG-I lectin. Although the overall fold of SasG-II is similar to that of SasG-I and Aap lectin domains, the local architecture in the vicinity of the glycan binding site differs in SasG-II. In particular, SasG-II lacks the characteristic aromatic residue (Aap Y580 or SasG-I W392) in the floor of the glycan binding pocket. These aromatic residues in Aap and SasG-I are presumed to form a stacking interaction with a bound glycan ligand, based on the loss of binding for the Y580A and W392A mutants of Aap and SasG-I, respectively ^16^. Thus, SasG-II may show a loss of glycan binding or a change in glycan specificity compared to Aap and SasG-I. Given their overall sequence and structural similarity, we conjectured that SasG-II might bind the same ligand as Aap and SasG-I as well as an additional ligand(s) not yet identified. Pre-incubating/blocking healthy human corneocytes with purified lectins from SasG-I and Aap was able to cross-inhibit SasG-II-mediated adhesion to corneocytes, and purified SasG-II was likewise able to cross-inhibit SasG-I and Aap-mediated adhesion. However, unlike the SasG-I and Aap lectins, which were found to bind both *N*-acetyl-D-lactosamine and 3’-sialyl-*N*-acetyllactosamine by Maciag et al. 2023 ^16^, the SasG-II lectin did not bind *N*-acetyl-D-lactosamine via ITC in our study. This was confirmed via corneocyte adhesion assays, where this purified glycan did not affect SasG-II-mediated adhesion to corneocytes. This suggests that SasG-II may bind the same glycan ligand as SasG-I and Aap, but may also bind elsewhere on the corneocyte receptor and occlude SasG-I and Aap from binding their respective glycan ligand.

AFM nanoimaging of corneocyte cell surfaces demonstrated weaker and less frequent adhesive interactions for SasG-I than for SasG-II, indicating that while both types strongly bind the ligand present on corneocytes, SasG-II may bind a broader variety of ligands and/or another glycan ligand on corneocytes. Additionally, pre-treatment of corneocytes with glycosidases resulted in different binding profiles between SasG-I and SasG-II. SasG-I-mediated adhesion was reduced by PNGase F, O-Glycosidase, α2-3,6,8 Neuraminidase, and the exoglycosidases α1-3,4 Fucosidase and β1-4 Galactosidase S, matching our previous report with the structurally similar *S. epidermidis* Aap ^15^. In contrast, SasG-II-mediated adhesion was reduced by only β-*N*- Acetylglucosaminidase S and β1-4 Galactosidase S. The only glycosidase that resulted in reduced adhesion of both SasG types was β1-4 Galactosidase S, suggesting that galactose may be an important terminal sugar residue in a shared ligand between SasG-I and SasG-II, while β-*N*-acetylglucosamine may be important in a ligand unique to SasG-II. Differentiated N/TERT keratinocytes treated with β-*N*-Acetylglucosaminidase S and β1-4 Galactosidase S further confirmed the corneocyte glycosidase data. Although SasG-I and Aap interact with complex N-glycan structures ^16^, SasG-II adhesion was unaffected by PNGase F, suggesting that SasG-II may bind to a distinct type of glycan. A likely candidate would be a core 2 O-glycan; these can contain terminal β1-4 galactose and β-*N*-acetylglucosamine ^38^ but are insensitive to cleavage by O-glycosidase ^39^. Interestingly, N-glycans containing lactosamine or sialyllactosamine and core 2 O-glycans are both prevalent species in the healthy skin glycome^35,36^.

While we do not know the exact structural nature of the ligand on the corneocyte receptor, our findings strongly suggest that the ligand(s) are expressed on desquamated, terminally differentiated keratinocytes and not basal keratinocytes, both SasG-I and SasG-II are likely to interact with a glycoprotein, and that SasG-II may also engage with a core 2 O-glycan structure (**Fig. 7**). The findings presented here as well provide further knowledge that could be used to therapeutically target and prevent *S. aureus* skin colonization for individuals at-risk for *S. aureus* infection in a prophylactic manner, potentially eliminating the need for traditional antibiotics that could contribute to further drug resistance in these individuals.

**Figure 7.**
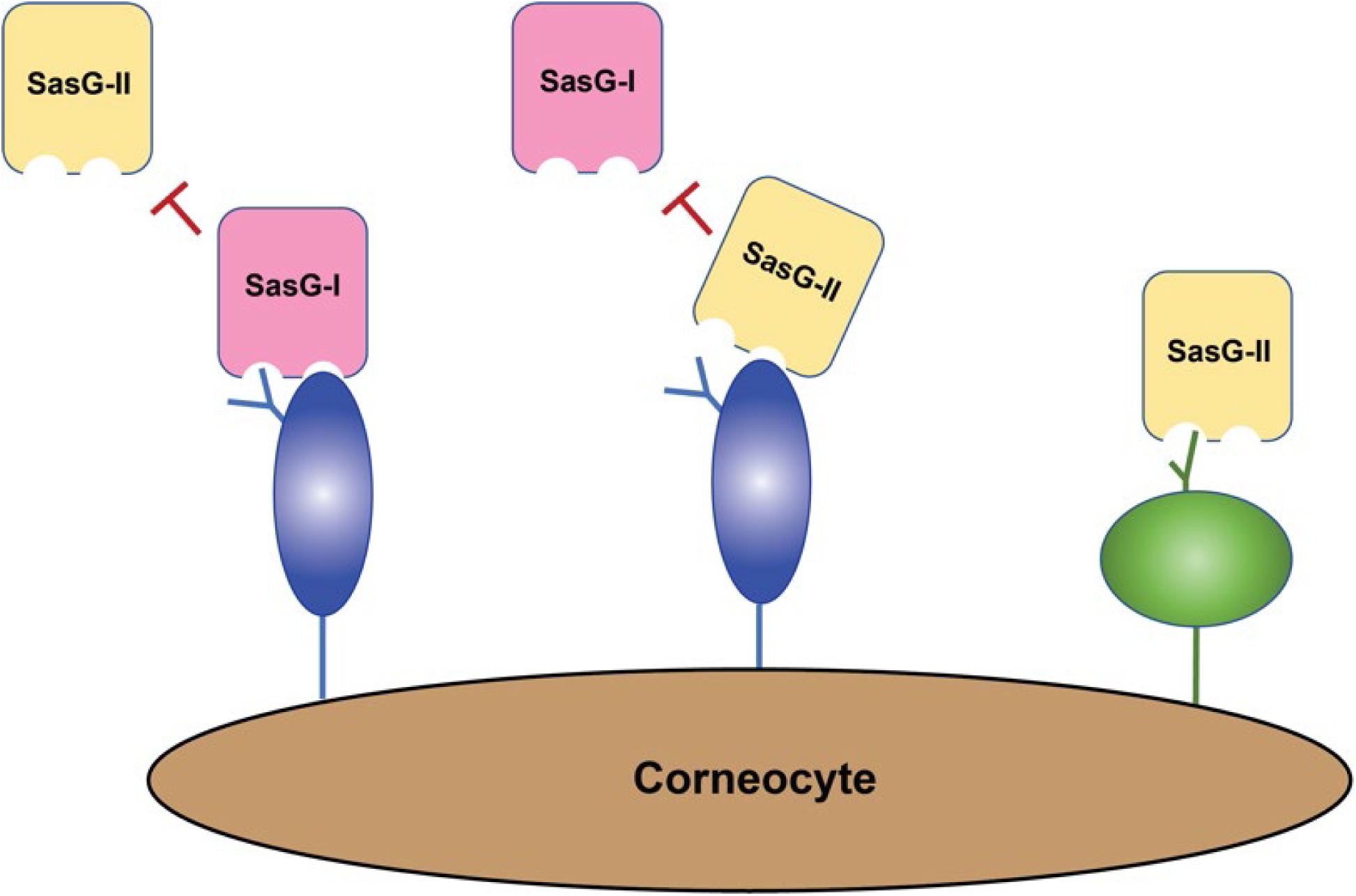
Model for SasG-I and SasG-II-mediated adhesion to healthy human skin corneocytes. SasG-I blocks adhesion of SasG-II, and likewise SasG-II can block adhesion of SasG-I, indicating they can all bind the same corneocyte receptor in some capacity. However, removal of N-glycans and core 1 or 3 O-glycans does not block SasG-II binding as it does with SasG-I, suggesting that SasG-II may bind a core 2 O-glycan structure elsewhere on the corneocyte receptor.

## Materials and methods

### SasG-I and SasG-II lectin alignment

The lectin protein sequences of SasG-I (from MSSA 502a) and SasG-II (from MRSA MW2) were aligned using Clustal Omega 1.2.4 ^46^. Sequence similarities, secondary structure elements, and relative accessibility were extracted with the Clustal Omega protein alignment and Type I/Type II lectin protein coordinates using the DSSP program in ESPript 3.0 ^47^.

### SasG phylogenetic analyses

Variation in SasG was analyzed in the context of *S. aureus* phylogenetic diversity. In brief, a previously curated data set of complete genomes from 574 *S. aureus* isolates, and 1 *S. argenteus* isolate that served as an outgroup, was obtained from the PATRIC database ^48–50^. PhyloPhlAn3 ^51^ was used to align conceptually translated protein sequences from the *S. aureus* proteome, curate the alignment, and perform phylogenetic analysis using default settings. Clonal complexes were identified on the phylogeny as clusters of related multilocus sequence types. SasG sequences were extracted from these proteomes and strains were sorted into three groups based on the state of their SasG sequence as full-length, truncated, or absent. Full-length SasG sequences were aligned with MUSCLE v3.8.31^52,53^ and clustered with CD-HIT v4.8.1 ^54,55^ at 100% identity. Gblocks v0.91b ^56^ was used with stringent settings (min. seq. for flank pos.: 85%, max. contig. nonconserved pos.: 4, min. block length: 10, no gaps in final blocks) to remove alignment gaps, including the SasG B-repeat sequences. PhyML ^57^ was used with the WAG model to infer a SasG phylogeny and to define two SasG allelic types. Mesquite v3.70 ^58^ was used to perform parsimony analysis with the *S. aureus* phylogeny to reconstruct SasG presence/absence and SasG allelic type; since the parsimony analysis ignores branch lengths it was not necessary to account for recombination events on the *S. aureus* phylogeny for this analysis. The R statistics package was used to perform a chi-squared goodness-of-fit test of the B-repeat distributions of the two SasG allelic types.

### SasG-II lectin crystallography

The SasG-II lectin domain was expressed as a fusion protein with an N-terminal hexahistidine tag followed by a tobacco etch virus (TEV) protease site using the pDest17-His plasmid in *E. coli* BLR (DE3) cells (Novagen). Protein expression and purification was conducted as described for SasG-I lectin domain ^16^. Crystals were grown via hanging drop diffusion. Protein stocks were at a concentration of 10 mg/mL in 20 mM Tris-HCl (pH 7.2) and 300 mM NaCl. Stocks were combined with mother liquor in equal parts (1 μL + 1 μL). Crystals appeared after five days in a condition of 100 mM HEPES (pH 7.8), 200 mM Ammonium Sulfate, 25%-33% BCS PEG SMEAR Medium (Molecular Dimensions, CalibreScientific). Cryoprotectant consisted of 80% mother liquor and 20% MPD; crystals were plunged into cryoprotectant prior to being flash frozen in liquid nitrogen. Data collection occurred at the Advanced Photon Source at Argonne National Lab through the Northeastern Collaborative Access Team (NE-CAT) on the 24-ID-E beamline. Data sets were collected at an oscillation range of 0.2°, collecting 900 frames, at a resolution range of 78.074-1.799 Å. Data indexing, space group assignment, scaling, and integration were carried out in the CCP4i suite^59^. Structure determination was carried out by PHENIX: PHASER using Aap lectin ^16^ as a search model for molecular replacement and building the initial SasG-II model with PHENIX: Autobuild. Iterative cycles of refinement and model building using data to a maximum resolution of 1.88 Å were carried out using PHENIX ^60^ and Coot ^61^, respectively. The final refined model was submitted for validation using MolProbity ^62^. Data collection and refinement statistics for the SasG-II lectin domain can be found in **Table S1**.

### Multiparametric imaging using single bacterial probes

SasG-I-expressing *S. carnosus*- SasG_COL_, SasG-II-expressing *S. carnosus*-*sasG*_MW2_, or *S. carnosus*-pALC2073 (EV) and a healthy skin corneocyte (0.5 cm × 0.5 cm) were immobilized on two different and separate areas on the bottom of a petri dish. The corneocyte was attached to one side of the petri dish using a double-sided transparent tape. 50 μL of diluted bacterial suspension in phosphate-buffered saline (PBS) was deposited on the other side of a Petri dish and allowed to adhere for 15 minutes at room temperature. The Petri dish was then carefully washed twice with PBS to remove non-adhering cells, after which 3 mL of PBS buffer was added to perform atomic force microscopy (AFM) experiments.

Single-cell probes were obtained by attaching a single bacterium to a colloidal probe. Colloidal probes were prepared as elucidated previously ^63^. Colloidal probe cantilevers were immersed for 60 minutes in Tris-buffered saline (TBS; Tris, 50 mM; NaCl, 150 mM; pH 8.5) containing 4 mg mL^-1^ of dopamine hydrochloride (Sigma-Aldrich), rinsed in TBS, and used directly for cell probe preparation. The nominal spring constant of the colloidal probe cantilever was determined by the thermal noise method ^64^, giving an average value of ∼0.06 N/m. The colloidal probe was brought into contact with a single isolated bacterium to catch it via electrostatic interaction with polydopamine and then moved on top of the corneocyte (kindly provided by Prof. Joan Geoghegan); proper attachment of the cell on the colloidal probe was checked using optical microscopy.

Multiparametric images of corneocytes were recorded in PBS using a bacterial probe under the Quantitative Imaging^TM^ mode available on the Nanowizard III and IV AFM (JPK Instruments, Germany). Images were obtained using a *S. carnosus* cell probe on top of a corneocyte at a scan area of 45 µm × 45 μm (256 pixels × 256 pixels), with an applied force of 0.5 nN, and a constant approach and retraction speed of 40 µm s^-1^ (z-range of 1 mm). For each condition, experiments were repeated for at least 3 different cell pairs.

### Isothermal titration calorimetry

ITC experiments were performed as previously described ^16^ using a MicroCal VP-ITC microcalorimeter. Analysis and fitting were done with ORIGIN software. Sample cells contained 20 μM of lectin protein (1.5 mL) and the syringe contained 1 mM of glycan (450 μl). The heat response from twenty glycan injections was measured; the first injection was a volume of 2 μl and the subsequent nineteen injections were at a volume of 14 μl each.

### Corneocyte collection

Desquamated corneocytes from the lower or upper arm near the elbow were collected from healthy human volunteers, as described in Mills et al. 2022 ^17^. Clear, adhesive tape stripping discs (d-Squame D100; Clinical & Derm) were used to collect corneocytes following cleaning and air-drying of the collection area with an alcohol wipe.

### Preparation of bacterial strains for corneocyte and N/TERT adhesion assays

The bacterial strains used in this study are listed in **Table S2**. Bacterial cultures were prepared as described in Mills et al. 2022 ^17^. All bacterial strains expressed superfolder green fluorescent protein (sGFP), either on plasmid pCM29 for *S. aureus* and *S. epidermidis* strains or chromosomally for *S. carnosus* strains. Strains were grown overnight at 37 °C with shaking aeration at 220 rpm in tryptic soy broth (TSB) in the presence of 10 µg/mL of chloramphenicol for plasmid maintenance. Strains were then sub-cultured at a 1:50 dilution in TSB with chloramphenicol and grown to an OD600 of ∼0.75. Strains were diluted to a final OD600 of 0.15 (∼10^7^ colony-forming units [CFU]/mL) after washing once in PBS at a 1:1 ratio. An OD600 of 0.15 was chosen to allow for adequate cell enumeration without clumping of bacterial cells ^15,17^.

### Corneocyte adhesion assays

The corneocyte adhesion assays were performed as described in Mills et al. 2022 ^17^.

#### Co-incubation with purified glycans

SasG-II-expressing *S. carnosus*-*sasG*_MW2_ was tested for adhesion to corneocytes following co-incubation with the purified glycans *N*- acetyl-D-lactosamine (Sigma-Aldrich) or 3’-sialyl-*N*-acetyllactosamine (Sigma-Aldrich). *N*-acetyl-D-lactosamine was prepared in PBS to a concentration of 1000 µM and diluted 2-fold from 1000 µM to 62.5 µM. 3’-Sialyl-*N*-acetyllactosamine was prepared in PBS to a concentration of 100 µM and diluted 2-fold from 100 µM to 6.25 µM. 300 µL of prepared glycans and 300 µL of prepared bacterial strains were co-incubated at room temperature for 20 minutes. The entire 600 µL was then incubated on corneocytes for 45 minutes at 37 °C. *S. carnosus*-pALC2073 was used as a negative control.

#### Pre-incubation with glycosidases

*S. carnosus* strains expressing SasG-I (*S. carnosus-sasG*_COL_) and SasG-II (*S. carnosus-sasG*_MW2_) were tested for adhesion to corneocytes following deglycosylation. Corneocytes were incubated with one of the following glycosidases for 24 hours at 37 °C in a moist chamber: PNGase F, O-Glycosidase, α1-2,3,6 Mannosidase, α1-3,4 Fucosidase, β-*N*-Acetylglucosaminidase S, β1-4 Galactosidase S, and α2-3,6,8 Neuraminidase (NEB). A 300 µL total solution for each enzyme contained: 3 µL enzyme, 30 µL GlycoBuffer (specific to each enzyme), and 30 µL BSA supplement for α1-3,4 Fucosidase or 30 µL zinc supplement for α1-2,3,6 Mannosidase, and addition of PBS up to 300 µL. After 24 hours, corneocytes were washed with PBS and subsequently incubated with the specified strains for 45 minutes at 37 °C. Each of the prepared solutions without the enzymes were used as control conditions for each experimental group.

#### Pre-incubation/blocking with purified lectins

Strains with SasG-II, *S. aureus* MW2 and *S. carnosus*-SasG_MW2_, were tested for adhesion to corneocytes following pre-incubation/blocking with purified recombinant lectins from Aap, Aap ΔY580A, SasG-I, SasG-I ΔW392A, and SasG-II. Protein purification of the lectins is described in Maciag et al. 2023 ^16^. Lectins were prepared to a concentration of 5 µM in 300 µL PBS. The prepared lectins were incubated on corneocytes at room temperature for 20 minutes, followed by incubation with 300 µL prepared bacterial strains for 45 minutes at 37 °C. *S. aureus* Δ*mgrA* Δ*sasG* or *S. carnosus*-pALC2073 (EV) were used as negative controls.

#### Pre-incubation/blocking with purified Type II full-length SasG and SasG-II A-domain

This assay followed a similar procedure as described in Mills et al. 2022 ^17^. *S. epidermidis* and SasG-I-expressing *S. carnosus*-*sasG*_COL_ were tested for adhesion to corneocytes following pre-incubation with purified full-length SasG-II or purified A-domain from *S. aureus* MW2. 300 µL of purified protein were prepared in PBS at a concentration of 100 µg/mL and incubated on corneocytes for 45 minutes at room temperature, followed by incubation with 300 µL prepared bacterial strains at 37 °C for 45 minutes. *S. epidermidis* Δ*ica* Δ*aap* or *S. carnosus*-pALC2073 (EV) were used as negative controls.

#### N/TERT adhesion assays

To study the adhesion of SasG to live host cells, immortalized N/TERT-2G keratinocytes were utilized ^65^. Here, low passage (<10) undifferentiated N/TERT-2G cells were seeded to 1×10^5^ cells/mL in either experiment media (EM) ^65^ or keratinocyte serum-free medium (KSFM, Gibco) in a 24-well tissue culture treated plate. For the assays on differentiated N/TERT-2G cells, the keratinocytes were both seeded in and allowed to differentiate in EM for 1 week with media changes every two days to promote the formation of a stratified and thick layer of epithelial cells. For assays using undifferentiated N/TERT-2G cells, cells were seeded in KSFM and were allowed form a 100% confluent monolayer (approximately 2-3 days growth) with daily media changes. All cells were grown at 37°C with 5% CO_2_.

On the day of experimentation, cells were inoculated with *S. carnosus*-pALC2073 (EV), *S. carnosus*-*sasG*_MW2_ (SasG-II), *S. carnosus*-*sasG*_COL_ (SasG-I), or *S. carnosus*-*sasG*_MW2ΔA_ prepared in PBS. The growth media was removed, and each strain was inoculated onto N/TERT-2G cells in technical triplicate at an MOI of 5 using a total inoculum volume of 100 uL. Upon addition of bacteria, the bacteria were allowed to incubate on the cells for 30 minutes following a 5-minute spin at 1000 rpm. After the 30-minute incubation, the N/TERT-2G cells were gently washed three times with 500 uL of sterile PBS per well to remove any un-adhered bacteria. 100 uL of 0.05% trypsin-EDTA (Sigma-Aldrich) was then added to each well and allowed to incubate for 10 minutes at 37°C with 5% CO_2_. The trypsinized cells were then resuspended in 400 uL sterile PBS, serially diluted, and plated to quantify the bacterial CFU/mL. Adhered bacterial cells were assessed as overall percent adhesion as well as percent cell association normalized to the *S. carnosus*-*sasG*_MW2_ input inoculum.

#### Pre-incubation with glycosidases

*S. carnosus* strains expressing SasG-I (*S. carnosus-sasG*_COL_) and SasG-II (*S. carnosus-sasG*_MW2_) were tested for adhesion to differentiated N/TERT-2G keratinocytes following deglycosylation. Cells were seeded and allowed to differentiate for 1 week as described above. Cells were incubated with glycosidases β-*N*-Acetylglucosaminidase S and β1-4 Galactosidase S at a 1:100 ratio in KSFM for 24 hours. After incubation, the cells were inoculated with the *S. carnosus* strains and followed the same adhesion and quantification protocol as described above.

## Acknowledgements

This work was funded by NIH/NIAID grant AI162964 to ARH, ABH, and PDF and <u>GM094363</u> to ABH. This research used resources of the Advanced Photon Source, a U.S. Department of Energy (DOE) Office of Science User Facility operated for the DOE Office of Science by Argonne National Laboratory under Contract No. DE-AC02-06CH11357. Aap diffraction data were collected at the Northeastern Collaborative Access Team (NE-CAT) beamline 24-ID-E, which is funded by the NIH through NIGMS (P30 GM124165). The Eiger 16M detector on 24-ID-E is funded by a NIH-ORIP HEI grant (S10OD021527).

