## Supplementary Materials for "*Staphylococcus aureus* skin colonization is mediated by SasG lectin variation"

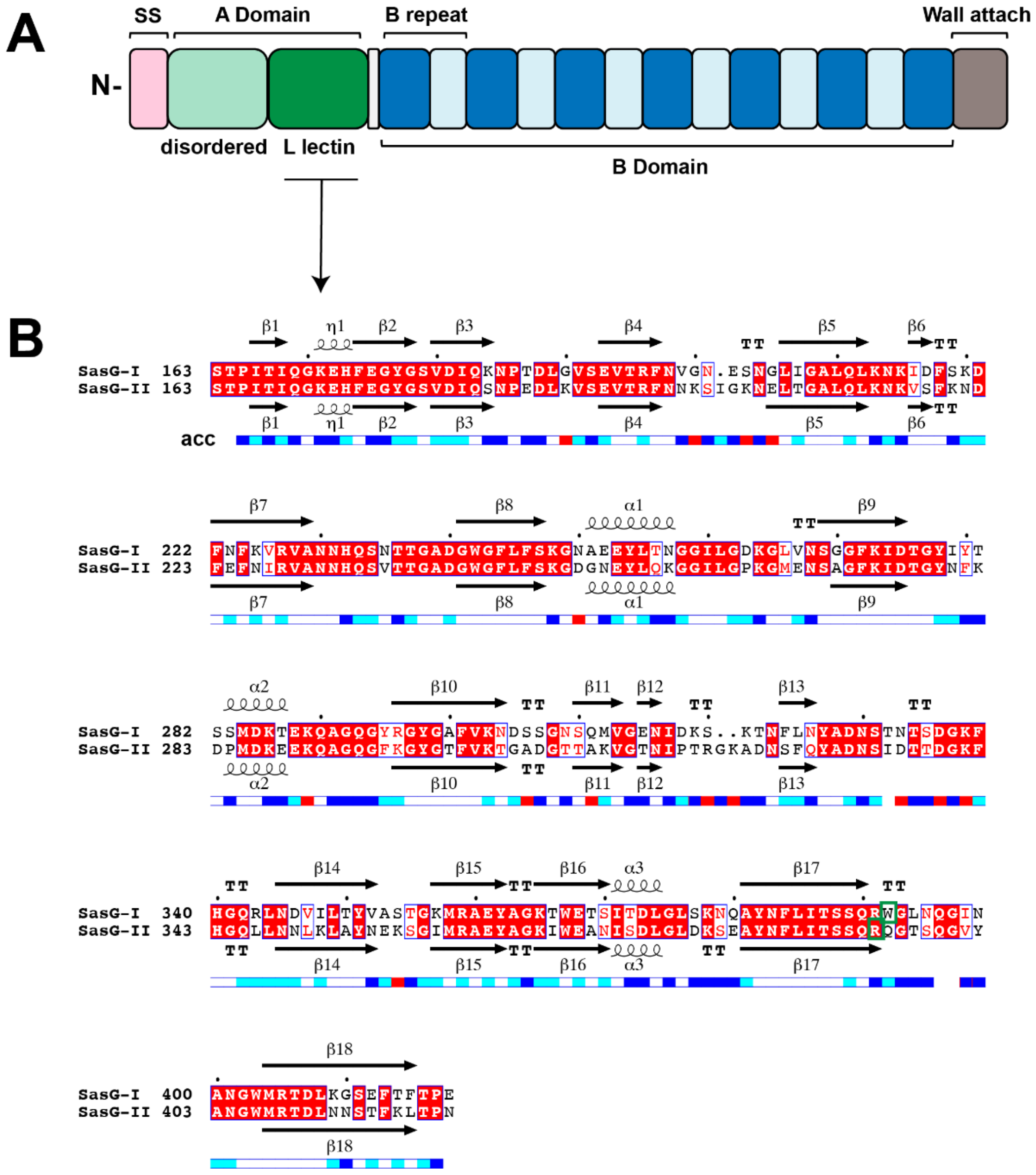


**Figure S1. The SasG A domain is variable in *S. aureus*. (A)** Schematic of the full-length SasG protein structure using MW2 SasG-II as a representative. Representative domains are shown, SS is signal sequence. (**B**) SasG-I and SasG-II lectin alignment. Sequence alignments from the SasG lectins of 502a (SasG-I) and MW2 (SasG-II) were performed using Clustal Omega. Predicted $\alpha$-helical, $\beta$-sheet, and turn (TT) structure elements were extracted from the structure coordinates of the SasG-I and SasG-II lectins using the DSSP program in ESPript. Sequences are listed next to SasG-I and SasG-II, where red boxes with white characters represent strict identity, red characters represent similarity in a group (in-Group Score), and blue frames represent similarity across groups (total score). Secondary structure elements from the SasG-I lectin protein coordinates are represented on the top of the alignment, and SasG-II is represented at the bottom of the alignment. The bottom configuration of the alignment (acc) represents the relative accessibility of each residue, where blue is accessible, cyan is intermediate, white is buried, and red is where residues between the sequences do not match. The key residues W392 for SasG-I and R394 for SasG-II are highlighted with green frames.

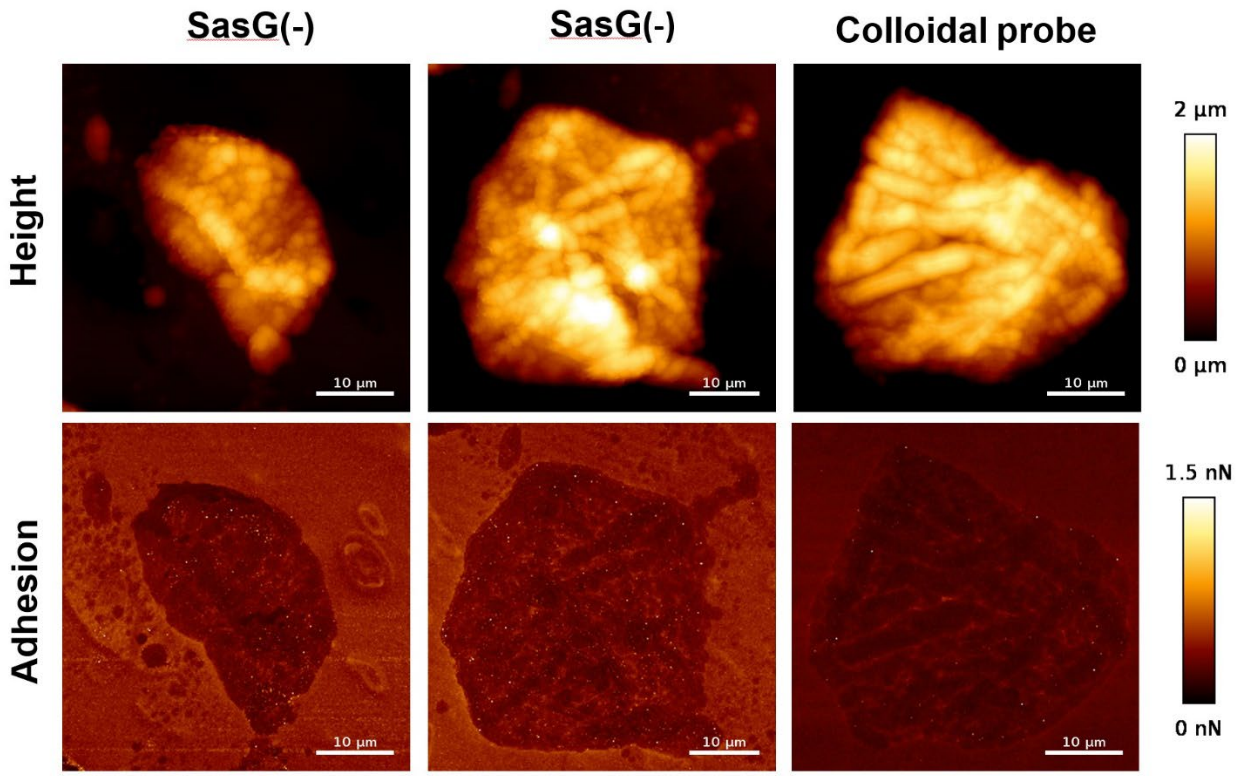


**Figure S2. Nanoscale multiparametric imaging of corneocytes using *S. carnosus*-pALC2073 EV (SasG[-]) or colloidal probes.**

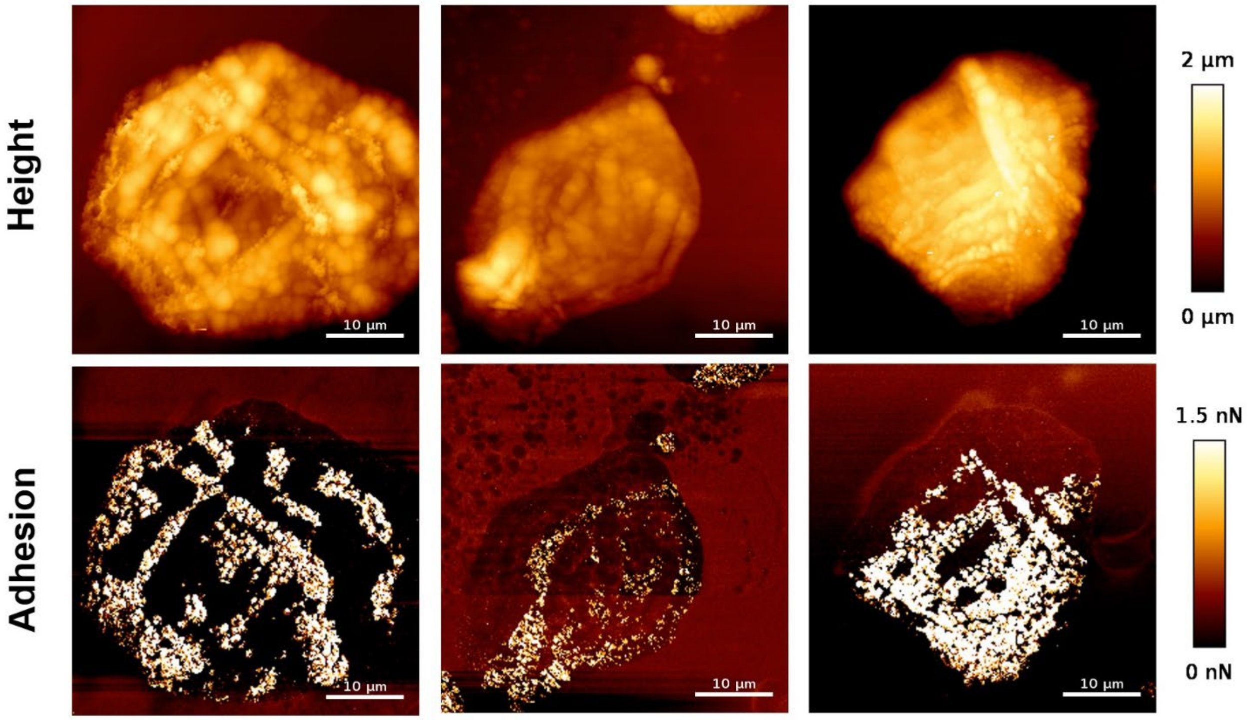


**Figure S3. Nanoscale multiparametric imaging of corneocytes using *S. carnosus*-*sasG*$_{MW2}$ (SasG-II) cell probes.**

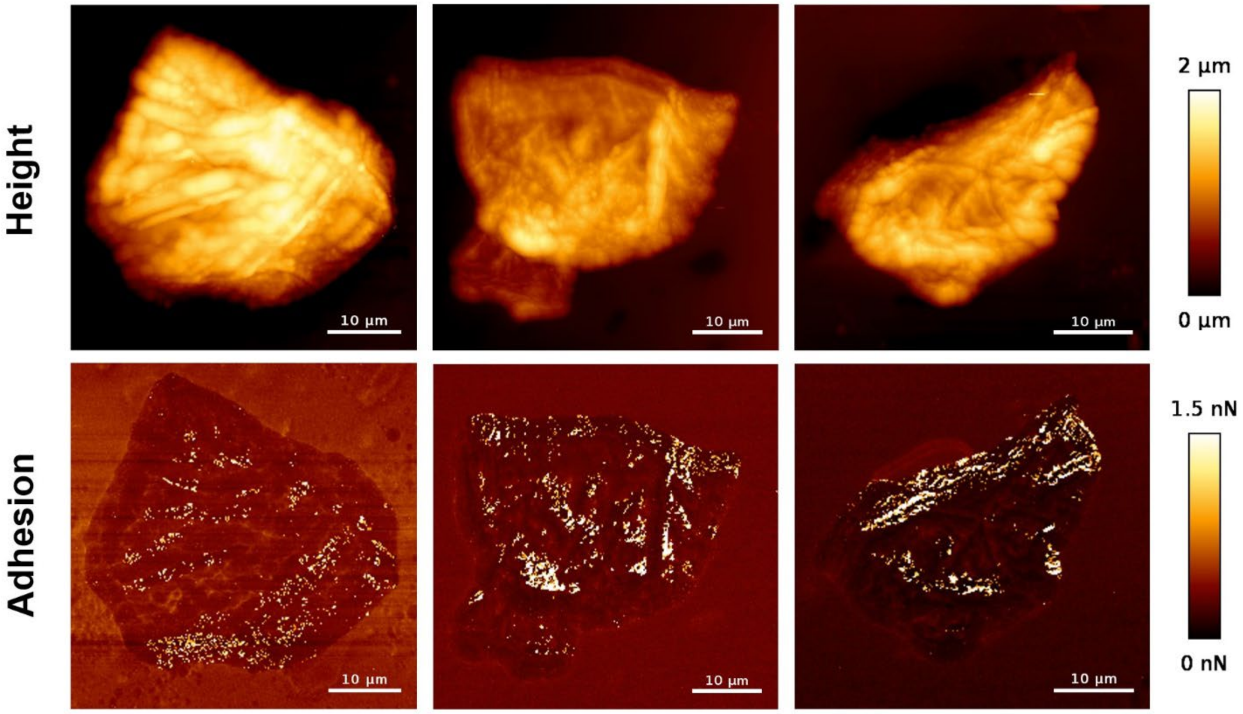


**Figure S4. Nanoscale multiparametric imaging of corneocytes using *S. carnosus*-*sasG*$_{COL}$ (SasG-I) cell probes.**

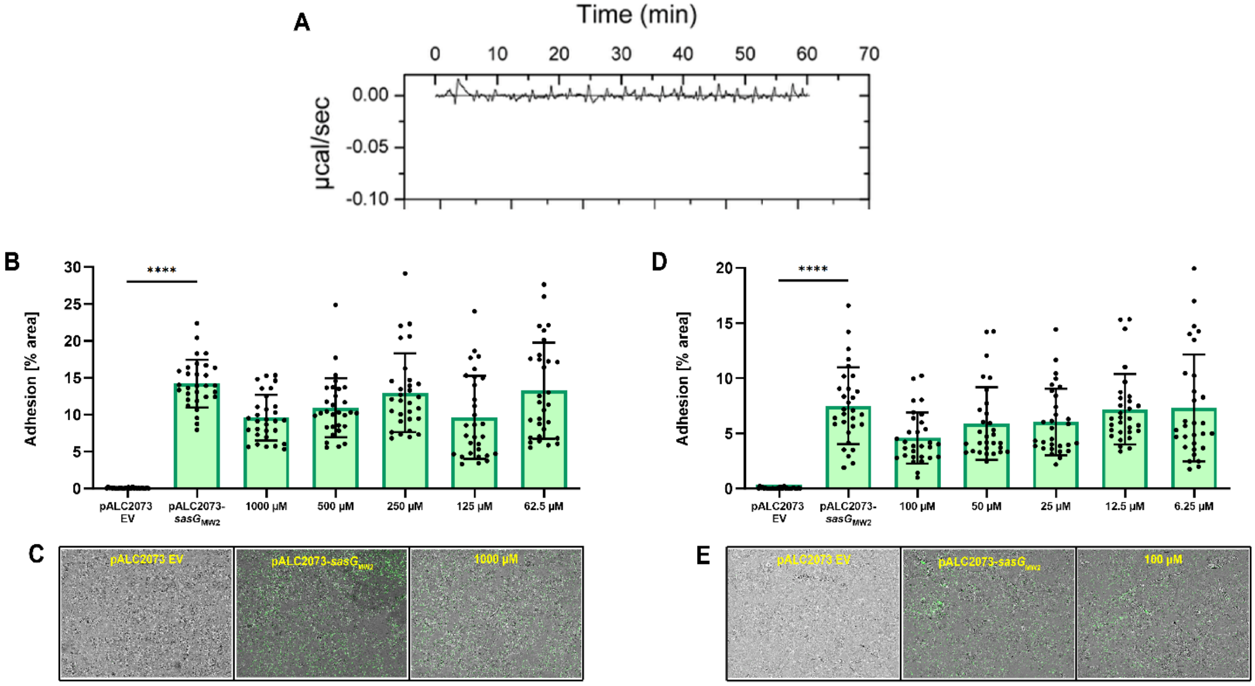


**Figure S5. Purified glycans do not affect Type II SasG-mediated adhesion to corneocytes.** (**A**) Type II SasG lectin did not bind to N-Acetyl-D-lactosamine via ITC. (**B**) Type II SasG-expressing *S. carnosus-sasG*$_{\mathrm{MW2}}$ was tested for adhesion to healthy human corneocytes after pre-incubation with 2-fold serial dilutions of N-Acetyl-D-lactosamine at a range of 1000 µM to 62.5 µM. None of the concentrations except 125 µM affected adhesion. (**C**) Representative bright-field (representing corneocytes) and green-channel (representing GFP-expressing bacteria) overlay microscopy images of experimental groups tested in Panel A. (**D**) Type II SasG-expressing *S. carnosus-sasG*$_{\mathrm{MW2}}$ was tested for adhesion to healthy human corneocytes after pre-incubation with 2-fold serial dilutions of 3'-Sialyl-N-acetyllactosamine at a range of 100 µM to 6.25 µM. None of the concentrations affected adhesion. (**E**) Representative bright-field (representing corneocytes) and green-channel (representing GFP-expressing bacteria) overlay microscopy images of experimental groups tested in Panel D. (**Panels B-E**) The percent area of adhesion in 10 images from three independent experiments (n = 30) was measured with Fiji ImageJ and analyzed in GraphPad Prism. Statistical significance was analyzed using the Kruskal-Wallis test (*****P*<0.0001).

**Table S1. Data collection and refinement statistics for SasG-II lectin domain.**

| | |
|---|---|
| Wavelength (Å) | 0.97933 |
| Resolution range (Å)[a] | 52.36–1.88 (1.947–1.88)[a] |
| Space group | P $2_12_12$ |
| Unit cell dimensions | |
| *a, b, c* (Å) | 78.60, 70.21, 43.54 |
| *α, β, γ* (°) | 90, 90, 90 |
| Total reflections | 40,304 (3,950) |
| Unique reflections | 20,226 (1,980) |
| Multiplicity | 2.0 (2.0) |
| Completeness (%) | 99.79 (99.85) |
| Mean I/sigma(I) | 5.81 (2.25) |
| Wilson B-factor | 20.27 |
| R-merge | 0.067 (0.297) |
| R-meas | 0.094 (0.421) |
| R-pim | 0.067 (0.297) |
| CC1/2 | 0.991 (0.780) |
| CC* | 0.998 (0.936) |
| Reflections used in refinement | 20,210 (1,977) |
| Reflections used for R-free | 1,999 (196) |
| R-work | 0.184 (0.250) |
| R-free | 0.253 (0.295) |
| CC (work) | 0.956 (0.894) |
| CC (free) | 0.914 (0.809) |
| Number of non-hydrogen atoms | 2052 |
| macromolecules | 1934 |
| ligands | 26 |
| solvent | 92 |
| Protein residues | 259 |
| RMS (bonds) | 0.006 |
| RMS (angles) | 1.15 |
| Average B-factor | 24.02 |
| macromolecules | 23.30 |
| ligands | 50.15 |
| solvent | 31.80 |
| Ramachandran favored (%) | 97.63 |
| Ramachandran allowed (%) | 2.37 |
| Ramachandran outliers (%) | 0.00 |
| Rotamer outliers (%) | 0.53 |
| Molprobity Clashscore (percentile) | 4.85 ($97^{th}$) |

[a] Statistics for the highest-resolution shell are shown in parentheses.

**Table S2. Bacterial strains used in this study.**

| Strain number | Species | Derivative Strain | Genotype | Plasmid |
|---|---|---|---|---|
| AH2983 | *S. epidermidis* | 1457 | Δ*ica* | pCM29 |
| AH2984 | *S. epidermidis* | 1457 | Δ*ica* Δ*aap* | pCM29 |
| AH4478 | MRSA | MW2 (USA400) | Δ*mgrA* | pCM29 |
| AH4834 | MRSA | MW2 (USA400) | Δ*mgrA* Δ*sasG* | pCM29 |
| AH5905 | *S. carnosus* | TM300 | Δ*SCA_2238*::sGFP Δ*SCA_2092* | pALC2073-*sasG*$_{MW2}$ |
| AH5907 | *S. carnosus* | TM300 | Δ*SCA_2238*::sGFP Δ*SCA_2092* | pALC2073 |
| AH5908 | *S. carnosus* | TM300 | Δ*SCA_2238*::sGFP Δ*SCA_2092* | pALC2073-*sasG*$_{COL}$ |
| AH6012 | *S. carnosus* | TM300 | Δ*SCA_2238*::sGFP Δ*SCA_2092* | pHC109 (pALC2073-*sasG*$_{MW2\Delta A}$) |
